# Coupling proximity biotinylation with genomic targeting to characterize locus-specific changes in chromatin environments

**DOI:** 10.1101/2024.07.26.605321

**Authors:** Pata-Eting Kougnassoukou Tchara, Jérémy Loehr, Jean-Philippe Lambert

## Abstract

Regulating gene expression involves significant and frequent changes in the chromatin environment at the locus level, especially at regulatory sequences. However, their modulation in response to pharmacological treatments or pathological conditions remain mostly undetermined. Here, we report versatile locus-specific proteomics tools to address this knowledge gap, which combine the targeting ability of the CRISPR/Cas9 system and the protein-labelling capability of the highly reactive biotin ligases TurboID (in CasTurbo) and UltraID (in CasUltra). CasTurbo and CasUltra enabled rapid chromatin protein labelling under mild conditions at repetitive sequences like centromeres and telomeres, as well as non-amplified genes. We applied CasUltra to A375 melanoma cell lines to decipher the protein environment of the *MYC* promoter and characterize the molecular effects of the bromodomain inhibitor JQ1, which targets bromodomain and extra-terminal (BET) proteins that regulate *MYC* expression. We quantified the consequences of BET protein displacement from the *MYC* promoter and found that it was associated with a considerable reorganisation of the chromatin composition. In addition, BET protein retention at the *MYC* promoter was consistent with a model of increased JQ1 resistance. Thus, through the combination of proximity biotinylation and CRISPR-Cas9-dependent genomic targeting, CasTurbo and CasUltra have successfully demonstrated their utility in profiling the proteome associated with a genomic locus in living cells.

**In Brief:** Kougnassoukou Tchara *et al*. report the development and application of CasTurbo and CasUltra, two locus-specific proteomics tools that fuse catalytically dead Cas9 to the engineered biotin ligases TurboID and UltraID. These tools enabled the quantitative mapping of locus-specific chromatin remodelling due to pharmacological inhibition.

**Highlights:**

- CasTurbo and CasUltra were developed for locus-specific label-free proteomics
- CasTurbo mapped the proteins localized to the centromeres and telomeres
- Proteins bound to the *MYC* promoter were quantified in melanoma cells with CasUltra
- CasUltra is compatible with investigating pharmacological treatment effects

**Graphical abstract:** 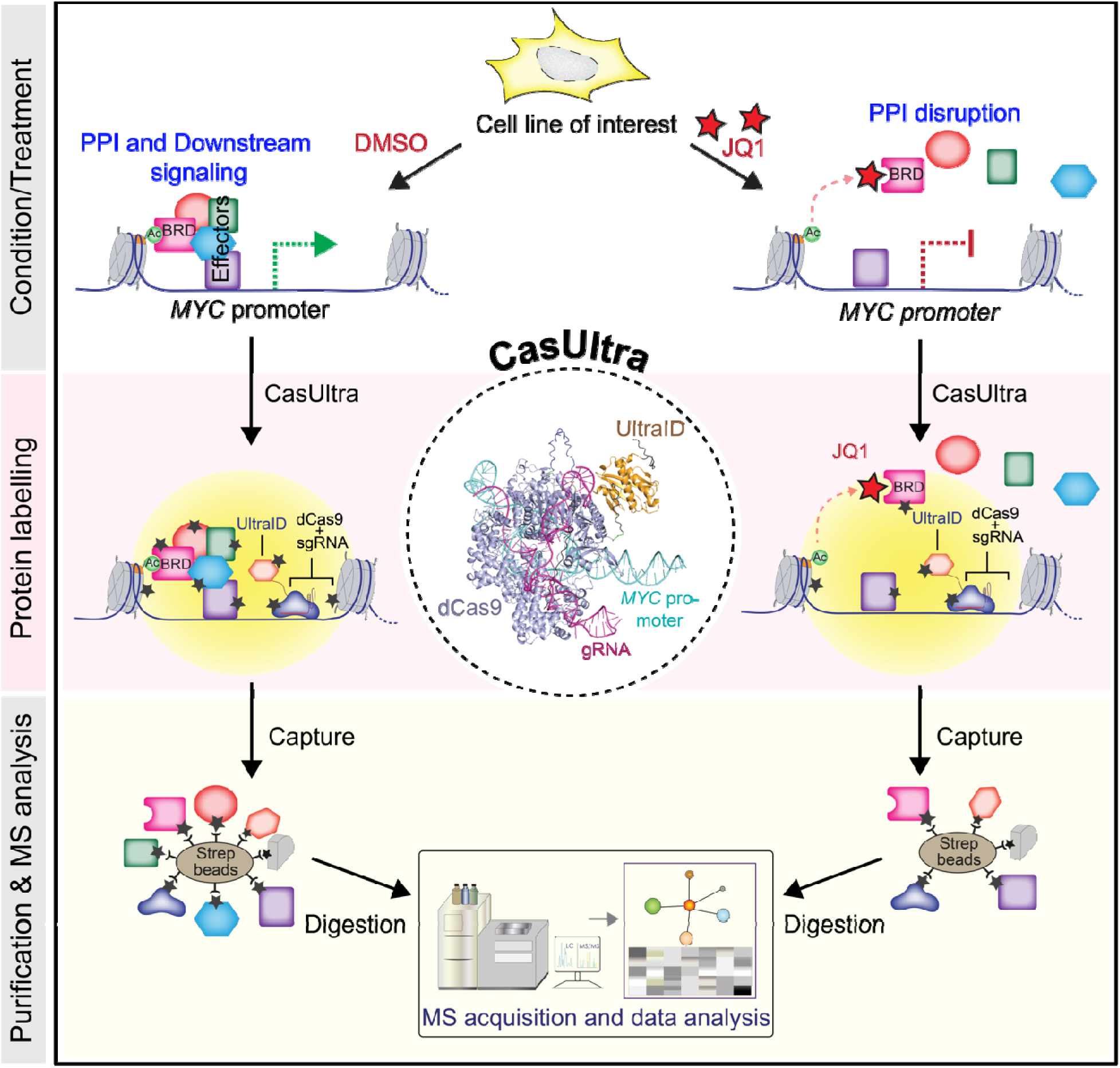

## Introduction

Lysine acetylation (Kac) is a post-translational modification found on thousands of proteins—most notably histones—and is the prominent recognition signal for bromodomains (BRDs) binding (1). Humans encode 61 distinct BRDs in 42 proteins. The BRD fold contains a large Kac binding pocket, and adjacent surfaces also contribute to Kac site specificity, providing numerous binding modalities for mono- and poly-acetylated substrates (2). The mainly hydrophobic binding cavity makes it well suited for the development of specific inhibitors that disrupt BRD-Kac associations (3–9). BRD-containing proteins are actively involved in numerous epigenetic processes. As Kac generally promotes transcription (10), many BRD-containing proteins do as well. However, recent research has revealed that the functions of BRD-containing proteins are more intricate than initially proposed and support signalling roles for many, with extensive cellular impacts (11).

The four human bromodomain and extraterminal (BET) proteins carry two distinct BRDs, bromodomain 1 (BD1) and bromodomain 2 (BD2), which display distinct functions and binding specificities for acetylated proteins depending on the context (**Figure 1A**) (12). BRD2 and BRD4 are commonly overexpressed in cancers (13–16), including melanoma (17–20). They are considered essential in most cancer cell lines in the Cancer Dependency Map (https://depmap.org/portal/) and are promising targets for clinical intervention (3–5, 8, 9, 21). We previously reported that BRD3 is functionally distinct from other BET proteins, as its expression is correlated with reduced proliferation and ribosomal RNA levels (22). The fourth BET protein, BRDT, is expressed only in the testes, where it is critical for spermatid maturation (23). BET proteins regulate transcription by organizing multiple protein complexes on specific promoters and enhancers (14, 24, 25). Disrupting BET BRD activity through chemical BET inhibition (BETi) drastically alters the transcriptomes of treated cells (26), presumably by disrupting these complexes (**Figure 1B**). Accordingly, over 50 clinical trials investigating them have been initiated to date (https://clinicaltrials.gov). However, whether modulating BRD protein function significantly alters the chromatin environment at specific loci remains to be quantitively defined.

**Fig. 1:**
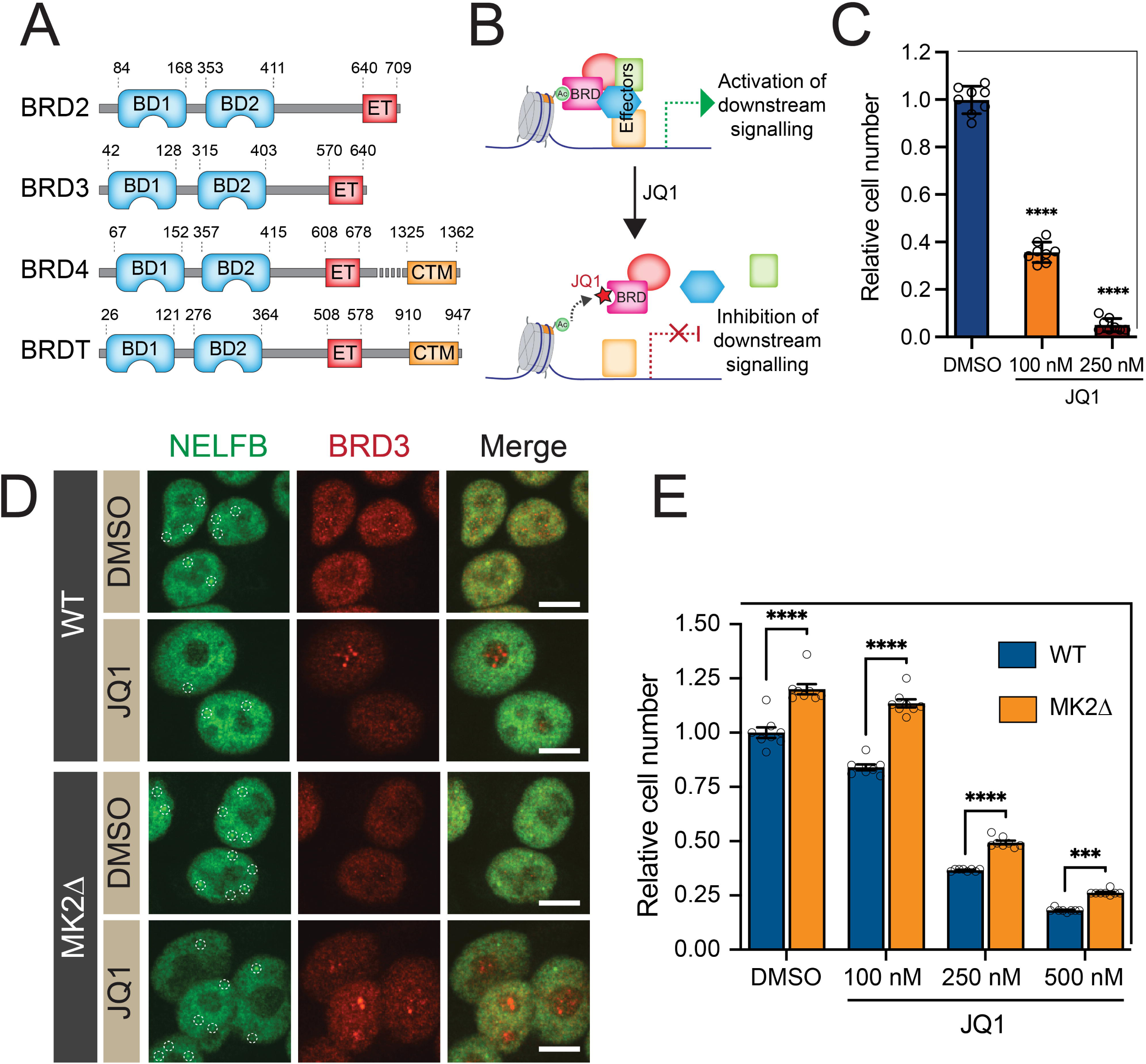
MK2 ablation and JQ1 treatment modulate chromatin composition. **A.** Domain organization of the BET proteins, with amino acid boundaries indicated. BD: bromodomain; ET: extra-terminal domain; CTM: C-terminal motif. Adapted from (22). **B.** Schematic of BET protein interactions with histone Kac marks and transcriptional regulators with and without JQ1. **C.** MTT-based proliferation assays of WT HeLa cells treated with increasing JQ1 concentrations for 8 d. The data are the mean ± the standard deviation (SD) of eight replicates per condition, and are representative of two independent experiments. **** *P* < 0.0001 compared to the DMSO sample, by *t*-test and one-way ANOVA. **D.** IF of NELFB and BRD3 in A375 lines (WT/MK2Δ) treated with DMSO or 500 nM JQ1 for 48 h. Scale bars: 10 µm. **E.** A375 WT and MK2Δ cell proliferation over 6 d in the presence of DMSO or different concentrations of JQ1. Cell density was quantified using a crystal violet assay. The data are the mean ± SD of eight replicates per condition, and are representative of two biological replicates. *** *P* < 0.001, **** *P* < 0.0001 compared to the DMSO sample, by *t*-test and one-way ANOVA.

The development of chromatin immunoprecipitation (ChIP) and its coupling to genome-wide measurement techniques has revolutionized our ability to study chromatin-associated proteins (27–30). ChIP sequencing (ChIP-Seq) and related techniques can precisely define how genetic or chemical perturbations change a protein’s chromatin localisation. A key feature of ChIP-Seq is the ability to generate in-depth data by amplifying tiny quantities of purified DNA. However, no such options exist for protein analysis, preventing the use of simple fractionation assays to quantify the chromatin at specific loci in detail (reviewed in (31)). In addition, the charged nature of chromatin can lead to hundreds of non-specific protein interactions during its biochemical isolation, particularly from S-phase cells (32). In recent years, direct adaptations of antibody-based ChIP protocols for proteomics analysis have provided new insights into chromatin composition (33–36), but not at the locus level. Although mini-chromosomes can enable significant amplification of a given locus (*e.g*., (37)), this approach is restricted to the few model organisms supporting their expression. The advent of efficient and flexible genome-targeting domains (zinc fingers, Cas9, *etc*.) has enabled mapping the chromatin environment at specific loci when combined with affinity purification coupled with mass spectrometry (AP–MS) (38). However, sensitivity remains an issue—as only proteins directly bound to the loci of interest can be biochemically purified in these assays, they are most effective for examining repetitive sequences (*e.g*., (39)). As we previously found that proximity-dependent biotinylation was complementary to AP–MS and allowed quantification of proteins with lower expression levels (40), we decided to adapt it for locus-specific chromatin mapping. While combining the genome targeting capabilities of catalytically dead Cas9 (dCas9) with a non-engineered abortive biotin ligase (BirA R118G; (41)) lacked the specificity to map chromatin environments because of its slow kinetics (42); CasID), the Carr (43), Sontheimer (44), and Liu (45) labs successfully fused dCas9 to the much faster APEX2 peroxidase (referred to herein as the CASPEX system).

Here, we adapted the CASPEX system to define how BETi remodels the chromatin environments surrounding key BET protein binding sites. We reengineered the system to incorporate the latest versions of the abortive biotin ligases TurboID (46) and UltraID (47), known as CasTurbo and CasUltra, respectively. As a proof of concept, we first benchmarked them at repetitive loci (centromeres and telomeres). Then, we used CasUltra to characterise molecular remodelling at the *MYC* promoter following BETi in melanoma cell lines. In doing so, we observed that the displacement of BET proteins from *MYC* was accompanied by a significant reorganisation of the chromatin composition, whereas the retention of BET proteins and the transcriptional machinery at the *MYC* promoter was associated with JQ1 resistance.

## Experimental procedures

### Plasmid construction

We used three dCas9-biotinylation enzyme (dCas9-BE) constructs with a N-terminal FLAG-tag (DYKDDDDK) and flanking triple SV40 nuclear localization sequences (3×NLS): 3×NLS-CASPEX (a derivative of the original CASPEX plasmid (Addgene #97421) generated by Myers *et al.* (43)), CasTurbo, and CasUltra. To generate the 3×NLS-CASPEX derivative (hereafter referred to as CASPEX for brevity), the original CASPEX plasmid and 3×FLAG-3×NLS_gBlock were digested with NcoI and SalI, then purified and ligated. CasTurbo was constructed by replacing the APEX2 sequence in CASPEX with TurboID. Briefly, APEX2 was excised from CASPEX by KpnI/AscI digestion. The resulting dCas9-containing vector was then ligated to an insert obtained by digesting gBlock_AscI-TurboID_KpnI with KpnI and AscI. The CasUltra plasmid was generated by removing TurboID from the CasTurbo plasmid by KpnI/AscI digestion and replaced with an UltraID insert digested from gBlock_AscI-CasUltraID_NotI, a gBlock fragment encoding the UltraID protein sequence reported by Kubitz *et al.* (47) (Addgene #172878). The single-guide RNAs (sgRNAs) used for site-specific targeting were cloned into a BsmBI-digested lentiGuide-Hygro-mTagBFP2 backbone (Addgene #99374). We generated sgRNA spacers by annealing the indicated complementary oligonucleotides as follows: 95°C (10 min), 85°C (1 min), 75°C (1 min), 65°C (1 min), 55°C (1 min), 45°C (1 min), 35°C (1 min), 25°C (1 min), with a final hold at 4°C. Annealed oligonucleotides were then ligated into a BsmBI-digested sgRNA lentiviral vector with T4 DNA ligase (New England Biolabs, #M0202L). The sgRNA-expressing vectors express blue fluorescent protein (BFP) as a visual indicator of sgRNA expression. Synthetic gBlock oligonucleotide and sgRNA sequences are provided in **Supplementary Table S1A**.

### Cell line production and growth

HeLa, HEK293T, and A375 cells and their subclones were cultured in Dulbecco’s Modified Eagle Medium (DMEM; Gibco, #11995-065) supplemented with 10% foetal bovine serum (FBS; Wisent) and antibiotics (penicillin/streptomycin; Gibco; LS15140122) in a humidified atmosphere containing 5% CO_2_. CASPEX, CasTurbo, and CasUltra plasmids were transfected into the cell lines with jetPRIME transfection agent (PolyPlus Transfection, #114-07) according to the manufacturer’s protocol. Puromycin (2 µg/mL) selection was initiated 72 h post-transfection and maintained for at least 2 weeks to generate dCas9-BE-expressing cell pools. We then performed limiting dilutions on the pools to select for clones stably expressing the dCas9-BE constructs. Isolated clones were subjected to immunofluorescence (IF) and WB to verify the doxycycline inducibility, proper subcellular localization, and biotinylation ability of their expressed dCas9-BE fusions. Screened clones were retained for lentiviral infection.

### CRISPR/Cas9-mediated MAPKAPK2 deletion in A375 and HeLa cells

The oligonucleotides used to generate CRISPR/Cas9-mediated *MAPKAPK2* knockouts (48) in A375 cells were designed using CRISPOR *via* the UCSD Genome Browser (49), as reported previously (50). Two guides targeting different exons were used to generate indels (see **Supplementary Table S1A** for oligonucleotide sequences). Oligonucleotides were annealed and ligated into Bbs1-digested PX459 V2.0 (Addgene #62988) with T4 DNA ligase (New England Biolabs, #M0202L). A375 cells were seeded in six-well plates, grown to approximately 40% confluency, and transfected with 2 µg of plasmid using jetPRIME transfection reagent (PolyPlus Transfection, #114-07) according to the manufacturer’s protocol. After 24 h, the cells were treated with 2 μg/mL Puromycin Dihydrochloride (Gibco, #A1113803) for 2 d. Then, the culture medium was replaced to allow the pool of cells to grow and limit plasmid integration into the genome. Guide efficiency was verified *via* tracking of indels by decomposition analysis (51). KO clones were then isolated by limiting dilution and screened for lack of MAPKAPK2 expression *via* western blot (WB). Genome editing events were confirmed *via* Sanger sequencing and deconvolution of complex DNA repair analysis (52).

### Lentivirus production

BFP- and sgRNA-encoding lentiviruses were produced by co-transfecting 2 µg of pCMV-VSV-G (Addgene, #8454), 3 µg of psPAX.2 (Addgene, #12260), and 3 µg of sgRNA plasmid into HEK293T cells seeded in 6-well plates at 70% confluence using jetPRIME transfection reagent according to the manufacturer’s protocol. The medium was replaced ≈15 h post-transfection, and lentiviruses were harvested 48 h later by centrifuging the cell medium at 500 × *g* for 5 min. The supernatant was filtered through a 0.45 μm filter into a new tube before transduction.

### Lentiviral infection

To express the sgRNAs, dCas9-BE-expressing clones were seeded in 6-well plates and grown until approximately 30% confluency (24–48 h) then treated with 8 µg/mL polybrene (Sigma, #107689) for 2–4 h. Cells were then transduced with 500 µL of freshly harvested lentivirus for 24–48 h and grown in medium containing 200 µg/mL Hygromycin B (Invitrogen, #10687010) for 2 weeks to select for cells that effectively incorporated the sgRNA. Three negative controls were generated for each cell line (no sgRNA, empty LentiGuide (further referred to as LentiG), and non-specific (NS) sgRNA-expressing), which were processed in parallel to the bait proteins. All subsequent experiments, except the proliferation assays, were performed on cells at approximately 70–80% confluency.

### Fluorescence-activated cell sorting (FACS)

Cells were seeded in 10 cm cell culture dishes, grown in DMEM supplemented with 10%FBS and antibiotics to 50% confluency, and induced with 1 µg/mL doxycycline for 24 h. After trypsinisation, filtration (0.45 μm), and washing with 1× PBS, 1 million cells co-expressing GFP (*i.e*., CasUltra) and BFP (*i.e*., an sgRNA) were sorted on a BD FACSAria Fusion Flow Cytometer and analysed using BD FACS Diva 8.0.3. BFP and GFP signals were detected at 450/50-nm and 530/30-nm, respectively. Cells highly expressing BFP were sorted unto two populations based on their GFP intensity (low and high). The sorted populations were expanded in DMEM supplemented with 10%FBS and antibiotics for downstream analysis.

### Chemical inhibitors

The bromodomain inhibitors (+)-JQ1 (Cayman Chemical, CAS#1268524-70-4), ABBV-075 (Cayman Chemical, Cat# 21033; CAS# 1445993-26-9), and ABBV-744 (Cayman Chemical, Cat# 30470; CAS# 2138861-99-9) were dissolved in dimethyl sulfoxide (DMSO) to obtain 50 mM stock solutions and then diluted to obtain 1–10 mM working solutions. DMSO was used as a vehicle control, and the working solution concentration for each treatment was chosen to minimise the inhibitor volume and reduce the intrinsic effects of DMSO (maximum 0.1% (v/v) of the medium). The cells were allowed to adhere to the culture dish for 2–24 h before BETi.

### Proliferation assays

Proliferation was assessed using the MTT cell proliferation assay (Roche, #11465007001) following the manufacturer’s procedure. Cells were seeded in a 96-well plate at 5000 cells/well, and the absorbance at 550–600 nm was measured using a Synergy H1 Hybrid Multi-Mode Reader (Agilent, Cat#11-120-531) at the indicated timepoints. Relative proliferation rates were determined by normalising values to the absorbance of the blank. For clonogenic assays, 10,000–20,000 cells were seeded in 12-well plates in DMEM supplemented with 10%FBS and antibiotics for 24 h before treatment with the appropriate drug. After the indicated incubation time, the medium was removed, and the cells were washed twice with 1× PBS, then stained with 200 µL of 0.5% crystal violet in 20% methanol (Sigma-Millipore, #1014080025, CAS #548-62-9). The plates were incubated at room temperature (RT) for 20–30 min, washed several times with 1× PBS, then imaged. Images were cropped using Adobe Illustrator. For absorbance measurements, crystal violet staining was performed as described above in a 96-well plate, and the dry samples were incubated with methanol (200 µL/well) for 30 min at RT in the dark under mild agitation. The absorbance at 570 nm was measured using a Synergy H1 Hybrid Multi-Mode Reader (Agilent, Cat#11-120-531). WT cells treated with DMSO were used as a reference for data normalization (as in (53)).

### Proximity-dependent biotinylation

Bait and control cell lines were processed in parallel. Briefly, cells were cultured in biotin-depleted medium (made by supplementing 500 mL of DMEM supplemented with 10%FBS and antibiotics with 1.6 mL of BioLock Biotin Blocking Solution (IBA LifeSciences, #2-0205-050), as first reported by Garloff *et al*. (54)) to reduce potential background due the high biotinylation activity of TurboID and UltraID. Cells were treated with DMSO or JQ1 for 48 h, with doxycycline (1 µg/mL) added 24 h into the treatment. The next day, the biotin-depleted medium was removed and the cells were washed twice with 1× PBS to remove all traces of BioLock then provided fresh DMEM supplemented with 10%FBS and antibiotics, as well as JQ1 or DMSO as needed, and incubated for 1 h (37°C, humidified atmosphere, 5% CO_2_) to allow for the cells to recover. Biotinylation was initiated by supplementing the medium with 500 µM biotin (Bio Basic Inc., Cat# BB0078) in DMSO, and the plates were incubated at 37°C for 20 min. The enzymatic reaction was stopped by removing the biotin-containing medium and washing the cells three times with cold 1× PBS before downstream treatment. For APEX2-mediated biotinylation, cells were first incubated for 30 min at 37°C with 500 µM biotin–phenol (Iris Biotech GmbH, #LS-3500.1000) in DMSO. Then, biotinylation was induced by the addition of 1 mM H_2_O_2_ (Sigma, #216763) to the medium and the cells were incubated at RT under horizontal shaking for 1 min. Subsequently, the medium was quickly removed, and the biotinylation reaction was quenched with three washes with ice-cold 1× PBS containing 10 mM sodium azide, 10 mM sodium ascorbate, and 5 mM Trolox, followed by three washes with ice-cold 1× PBS. Proximity-dependent biotinylation experiments were performed in triplicate, with each biological replicate sourced from two 150 mm plates, and analysed by MS. After biotinylation, the cells were harvested on ice using a scraper, pelleted by centrifugation at 1,000 × *g* at 4°C, and either immediately processed or snap-frozen and stored at –80°C until purification.

### Immunofluorescence

Cells were seeded in 12-well plates containing coverslips in complete media and grown for 48 h to reach 50-75 % confluency. After 24 h of induction with 1 µg/mL doxycycline, the cells were fixed with 4% paraformaldehyde in 1× PBS for 15 min at RT, and then stained for FLAG, biotin (with fluorophore-conjugated streptavidin), or CREST (ImmunoVision, #HCT0100). Image Z stacks at an interval of 0.15 µm were acquired using an inverted microscope (DMI 6000B, Leica) equipped with a confocal unit (CSU10, Yokogawa) at 63× magnification. Z stacks were deconvolved using Volocity (Quorum Technologies) and displayed as maximum intensity projections. Images were cropped using Adobe Photoshop. For all quantitative comparisons, we used identical imaging conditions (including exposure times).

### Western blotting

For WB analysis, 30 μg of protein lysate was resolved by SDS–PAGE, transferred to nitrocellulose, blocked in Tris-buffered saline containing 5 mg/mL non-fat milk and 1% Tween-20 for 1 h at RT under horizontal rotation, and incubated with primary and secondary antibodies as outlined in **Supplementary Table S1B**. Chemiluminescence generated using the using Clarity Western ECL Substrate (Bio-Rad; #1705061) was detected using either a ChemiDoc Imaging System (Bio-Rad, Cat # 12003153) or on film. Films were scanned, and WB figures were assembled using Adobe Photoshop and Adobe Illustrator.

### Chromatin immunoprecipitation

Cells were cultured to 50–60% confluency in 150 mm culture plates, doxycycline-induced for 24 h, and biotinylated as described above. All buffers—from cell lysis to immunoprecipitation—were supplemented with protease inhibitors (Sigma, #P8340) and phosphatase inhibitors (Sigma, # P5726 and # P0044) and handled on ice or at 4°C. Briefly, after biotinylation, cells were fixed with 1% (v/v) formaldehyde (Sigma, #252549) for 15 min at RT under mild agitation to stabilise the protein–DNA complexes by crosslinking, quenched with 125 mM glycine for 5 min, and washed three times with 15 mL of ice-cold 1× PBS. The cells were then scraped into 15 mL conical tubes in 5 mL PBS and centrifuged (1,000 × *g*) for 5 min at 4°C to separate the supernatant. The crosslinked cell pellet was resuspended (1 mL/2×10^7^ cells) in lysis buffer (50 mM Tris-HCl, pH 7.5; 1 mM EDTA, pH 8.0; 140 mM NaCl; 1% NP-40), incubated on ice for 10 min, and centrifuged (2,000 × *g*) for 10 min at 4°C. The supernatant was discarded, and the nuclei pellet was resuspended (0.5 mL/2×10^7^ cells) in ice-cold nuclear lysis buffer (50 mM Tris-HCl, pH 7.5; 2 mM EDTA, pH 8.0; 0.5% sodium deoxycholate; 1% SDS). The suspension was sonicated at 4°C with a Bioruptor (Diagenode, #B01020001) using conditions optimized to generate 200–500 bp chromatin fragments. The chromatin fraction was cleared by centrifugation (17,000 × *g*) for 10 min at 4°C, and the supernatant was collected into a fresh tube. Most of the sheared chromatin was snap-frozen and stored at –80°C until ChIP, but a 20 µL aliquot was processed to determine the DNA concentration and size distribution as follows. Crosslinks were reversed by incubation in 0.25 M NaCl at 65°C for 3–4 h or overnight, followed by treatment with 0.1 mg/mL Monarch RNase A (New England Biolabs, #T3018L) for 30 min at 37°C and then 0.5 mg/mL proteinase K (Invitrogen, #25530049) for 1 h at 55°C. Finally, the free DNA fragments were purified using a Monarch PCR & DNA Cleanup Kit (5 μg) (New England Biolabs, #T1030S) and quantified, and their size distribution was analysed using agarose gel electrophoresis. For ChIP, 100 µL of chromatin fragments were diluted in 900 µL dilution buffer (20 mM Tris-HCl, pH 8.0; 1 mM EDTA, pH 8.0; 0.1% sodium deoxycholate; 140 mM NaCl; 0.01% SDS; 1% NP-40) and incubated with 25 µL Dynabeads Protein A of Immunoprecipitation (Invitrogen, #10002D) pre-coupled with 2.5 µg of anti-FLAG antibody (Sigma, #F1804) or rabbit IgG (Sigma, #I8140) for 3 h at 4°C under gentle rotation. After chromatin capture for 5 min at RT with rotation, the beads were washed sequentially with 500 µL of low-salt buffer (20 mM Tris-HCl, pH 8.0; 2 mM EDTA, pH 8.0; 0.1% SDS; 1% Triton X-100; 150 mM NaCl), high-salt buffer (20 mM Tris-HCl, pH 8.0; 2 mM EDTA, pH 8.0; 0.1% SDS; 1% Triton X-100; 500 mM NaCl), LiCl buffer (20 mM Tris-HCl, pH 8.0; 2 mM EDTA, pH 8.0; 0.5% sodium deoxycholate; 0.1% NP-40; 250 mM LiCl), and Tris-EDTA buffer (10 mM Tris-HCl, pH 8.0; 1 mM EDTA). After the final wash, the beads were resuspended in 100 µL of elution buffer (1% SDS, 100 mM NaHCO_3_), rotated for 15 min at RT, and incubated at 65°C for 10 min; the supernatant was recovered into a fresh tube. For each sample, 10 µL of sheared chromatin was diluted 10× with elution buffer and used as an input sample. Both the immunoprecipitates and input samples were processed for crosslink reversal and DNA purification as described above and stored at –80°C until qPCR.

### qPCR

Sequence-specific PCR primers were designed according to Gao *et al.* (44). ChIP-qPCR input chromatin was used to generate standard curves for the amplification of each target primer set to validate their efficiency and determine the amount of DNA required for qPCR. Samples were diluted at a 1:5 ratio with H_2_O prior to qPCR using Luna Universal qPCR Master Mix (New England Biolabs, #M3003) according to the manufacturer’s instructions. Reactions were performed in 96-well EnduraPlate optical reaction plates (Applied Biosystems, #4483354) using a QuantStudio 3 Real-Time PCR System (Applied Biosystems, #A28567), with three technical replicates per experimental condition. The data were analysed using the fold enrichment method and normalized to the cycle threshold values of GAPDH, as described (55). The amplification targets were centromeres (alpha satellites), telomeres, and GAPDH. PCR products were analysed by agarose gel electrophoresis to verify their sizes. Primers used for qPCR are listed in **Supplementary table S1A**.

### Streptavidin purification

The streptavidin-based purification protocol was adapted from Lambert *et al.* (2019) (22) with a few modifications. Each sample consisted of a biotinylated cell pellet from two 150 mm plates freshly snap-frozen or stored at –80°C. Briefly, samples were thawed and resuspended in 1.5 mL ice-cold radioimmunoprecipitation assay (RIPA) buffer containing 50 mM Tris-HCl (pH 7.5), 150 mM NaCl, 1% NP-40, 1 mM EDTA, 1 mM EGTA, 0.1% SDS, and 0.5% sodium deoxcycholate. PMSF (1 mM), DTT (1 mM), and Protease Inhibitor Cocktail (P8340, 1:500, Sigma-Aldrich) were added immediately before use. The lysates were sonicated using a Sonic Dismembrator Series 60 (four 10 s bursts, 35% amplitude), treated with 1 µL Turbonuclease (Sigma, #T4330), and centrifuged (20,000 × *g*) for 20 min at 4°C. The supernatant was transferred into a new 2 mL tube. Streptavidin Sepharose (GE Healthcare, cat. 17-5113-01; 60 μL/sample) was pre-washed three times with 1 mL of RIPA lysis buffer by pelleting the beads with gentle centrifugation and aspirating off the supernatant before the next washing. Washed beads were added to each sample and the suspension was gently rotated for 3 h at 4°C to allow the biotinylated proteins to bind the streptavidin beads. The beads were gently pelleted and then washed twice with 1 mL RIPA buffer (without protease inhibitors) and three times with 1 mL of 50 mM ammonium bicarbonate (pH 8.0). After the final wash, the beads were pelleted, and the excess liquid was aspirated. The beads were re-suspended in 100 μL of 50 mM ammonium bicarbonate (pH 8.0) supplemented with 1 μg of trypsin and rotated overnight at 37°C. The next day, an additional 1 μg of trypsin was added, followed by further incubation for 2–4 h. The beads were pelleted, and the supernatant was transferred to a fresh tube. The beads were rinsed twice with 100 μL HPLC-grade water, and the wash fractions were combined with the supernatant. The digested samples were acidified with 50% formic acid to a final concentration of 2% and vacuum dried in a Speedvac. The dry peptides were then isopropanol-fractionated or stored at −80°C until further processing.

### Peptide fractionation

Peptides were fractionated as described by Deng *et al.*, with slight modifications (56). Dry peptides from digested streptavidin-purified digested proteins were solubilized in 50 µL of 95% acetonitrile (ACN). We used a 1:1 mixture of Sera-Mag Carboxylate-Modified Magnetic Beads (CMMBs; 65152105050250 and 45152105050250, GE Healthcare) to bind the solubilized peptides. Briefly, 50 µL of CMMBs were washed with 250 µL of 95% ACN, resuspended in 50 µL of 95% ACN, combined with the peptides, and horizontally shaken at 220 rpm (#Excella E25, New Brunswick Scientific) for 20 min at RT to allow the peptides to bind to the beads. The peptides were sequentially eluted from the CMMBs using 100 µL of eluent (90%, 80%, and then 70% isopropanol) by pipetting up and down 15 times, followed by a 10 min incubation under horizontal agitation at RT. The eluted peptide fractions were dried under vacuum centrifugation.

### Peptide cleanup

LC–MS-grade reagents were used, and 20 µL of each buffer was added. After adding a buffer or sample, the column was centrifuged for 30 s at 500 × *g* (for methanol and buffer B (80% ACN and 2% formic acid)), or for 3 min at 1,500 × *g* (for buffer A (H_2_O with 2% formic acid)). Briefly, a C_18_ StageTip column was activated with 100% methanol and sequentially conditioned with buffer B and buffer A. The accumulated flowthrough was discarded. The dried peptide eluate was solubilized in 20 µL of buffer A, vigorously vortexed, left to stand for at least 5 min at RT, and centrifuged (1,500 × *g*) for 30 s. The supernatant was applied to the column. After centrifugation, the flowthrough was reloaded and centrifuged again to maximize peptide binding. The column was then washed twice with buffer A before three elutions with buffer B into a new tube. The eluted sample was dried by vacuum centrifugation and stored at –80°C until LC–MS.

### Experimental design and statistical rationale for MS experiments

For each MS analysis, three independent biological replicates of each bait were processed. Negative controls were processed in each batch of samples. The order of sample acquisition on the LC–MS/MS system was randomized.

### Protein identification by MS

Peptide samples were separated using online reversed-phase nanoscale capillary LC on a Dionex UltiMate 3000 nanoRSLC chromatography system (Thermo Fisher Scientific) and analysed by electrospray ionization tandem mass spectrometry (MS) on an Orbitrap Fusion mass spectrometer (Thermo Fisher Scientific) equipped with a nanoelectrospray ion source. Peptides were trapped at 20 μL/min in a loading solvent (2% ACN, 0.05% trifluoroacetic acid) on a 5 mm × 300 μm C_18_ PepMap cartridge pre-column (Thermo Fisher Scientific) for 5 min. Then, the pre-column was switched online with a self-made 50 cm × 75 μm internal diameter separation column packed with ReproSil-Pur C_18_-AQ 3-μm resin (Dr. Maisch HPLC), and the peptides were eluted with a linear gradient from 5–40% solvent B (A: 0.1% formic acid; B: 80% ACN, 0.1% formic acid) for 90 min at 300 nL/min. Mass spectra were acquired using the data-dependent acquisition mode in Thermo XCalibur software (version 3.0.63). Full-scan mass spectra (350–1800 m/z) were acquired in the Orbitrap using an automatic gain control target of 4e5, a maximum injection time of 50 ms, and a resolution of 120,000. The spectrometer was internally calibrated by locking onto the mass of the siloxane ion (m/z 445.12003). Each MS scan was followed by acquisition of the fragmentation spectra of the most intense ions for a total cycle time of 3 s (top-speed mode). The selected ions were isolated using the quadrupole analyser in a 1.6 m/z window and fragmented by high-energy collision-induced dissociation with 35% collision energy. The resulting fragments were detected by the linear ion trap at a rapid scan rate with an automatic gain control target of 1e4 and maximum injection time of 50 ms. Dynamic exclusion of previously fragmented peptides was set for a period of 20 s and a tolerance of 10 ppm.

### Protein identification and quantification

MaxQuant (version 2.0.3.0; http://www.maxquant.org/) was used to identify and quantify proteins in the dataset, using default parameters with a few exceptions. Peptides were identified from MS/MS spectra using UniProt Human Proteome UP000005640_9606 (UniProt release 2023_03, containing 20,586 proteins) as a reference database. Fractions from the same sample were set as individual experiments in the same parameter group. Replicates were processed independently. Methionine oxidation and N-terminal acetylation were set as variable modifications. Trypsin was selected as the protease, with up to two missed cleavages allowed. The peptide length range was set from 7–25, with a maximum mass of 4600 Da. The false discovery rate was set to 0.01 for peptides, proteins, and sites. The analysis was performed using label-free quantification with MaxLFQ implemented in MaxQuant (57). The “proteinGroups” .txt files produced by MaxQuant were used as input files to analyse and visualise the data in volcano plots with LFQ-Analyst using default settings (58). All materials used in this study are summarized in **Supplementary Table S1B**. Detailed MS data are provided in **Supplementary Table S2**.

### MS data archiving

All MS files used in this study were deposited to MassIVE (http://massive.ucsd.edu) and assigned MassIVE number MSV000094950. The files can be downloaded at ftp://MSV000094950@massive.ucsd.edu using “chromatin” as the password prior to publication.

### Statistical analysis

All schematic overview figures were created using Adobe Illustrator (v26.0.2, Adobe). Prism 10 (GraphPad Software) was used for all statistical analyses and figure creation. At least two independent biological replicates were performed for each experiment. Two-tailed (paired/unpaired) student’s *t*-test and two-way ANOVA were used to analyse the significance of differences between two groups and among several groups, respectively. *P* < 0.05 was considered significant.

## Results

### The BET protein inhibitor JQ1 reduces cell proliferation and disturbs the chromatin proteome

The antiproliferative effects of BETs inhibitors, notably JQ1, have been widely documented by us and others (4, 59–62). By lodging itself in the hydrophobic pocket of the bromodomain, JQ1 abrogates the interaction between BET proteins and their acetylated interaction partners, leading to alterations in downstream chromatin signalling (**Figure 1B**). This includes inhibiting proliferation (**Figure 1C**), relocalising BRD3 to the nucleolus, and increasing its interaction with nucleolar partners following JQ1 treatment (**Figure 1D**) (22). Moreover, JQ1 significantly reduces chromatin foci containing negative elongation factor B (NELFB) (50), a subunit of the NELF complex, which, along with the DRB sensitivity-inducing factor complex, pauses transcriptional elongation by RNA polymerase II after its initial burst. As we recently reported (50), as observed following JQ1 treatment, genetic ablation of MAPK activated protein kinase 2 (MAPKAPK2; hereafter referred to as MK2) in melanoma cell lines significantly enhanced both JQ1 resistance and the number of NELFB foci (**Figure 1D-E**), indicating intense transcription (63). Therefore, we hypothesized that BETi drastically remodels the chromatin composition and sought to precisely map these changes at key genes.

### Generation and characterization of dCas9-fused biotinylation enzymes (dCas9-BE models)

To profile chromatin proteome reorganization at the locus level, we created the CasTurbo and CasUltra tools. Briefly, we coupled the genome-targeting capabilities of dCas9 with the interactome-profiling capacities of the biotinylation enzymes (BEs) TurboID (CasTurbo) and UltraID (CasUltra), and compared them to the previously reported APEX2-based tool (CASPEX) (43). The cell lines of interest were first transfected with a plasmid encoding a doxycycline-inducible dCas9-BE fusion (dCas9-BE; **Figure 2A**, left) and puromycin-selected (**Figure 2B**). Pools of stably expressing dCas9-BE cells were characterized by IF and WB, and clonal cell lines were prepared by limiting dilution. Optimal clones stably expressing each dCas9-BE (based on their nuclear localization, mild expression levels, and biotinylation abilities) were then lentivirally transduced with sgRNA (**Figure 2A**, right) and hygromycin-selected.

**Fig. 2:**
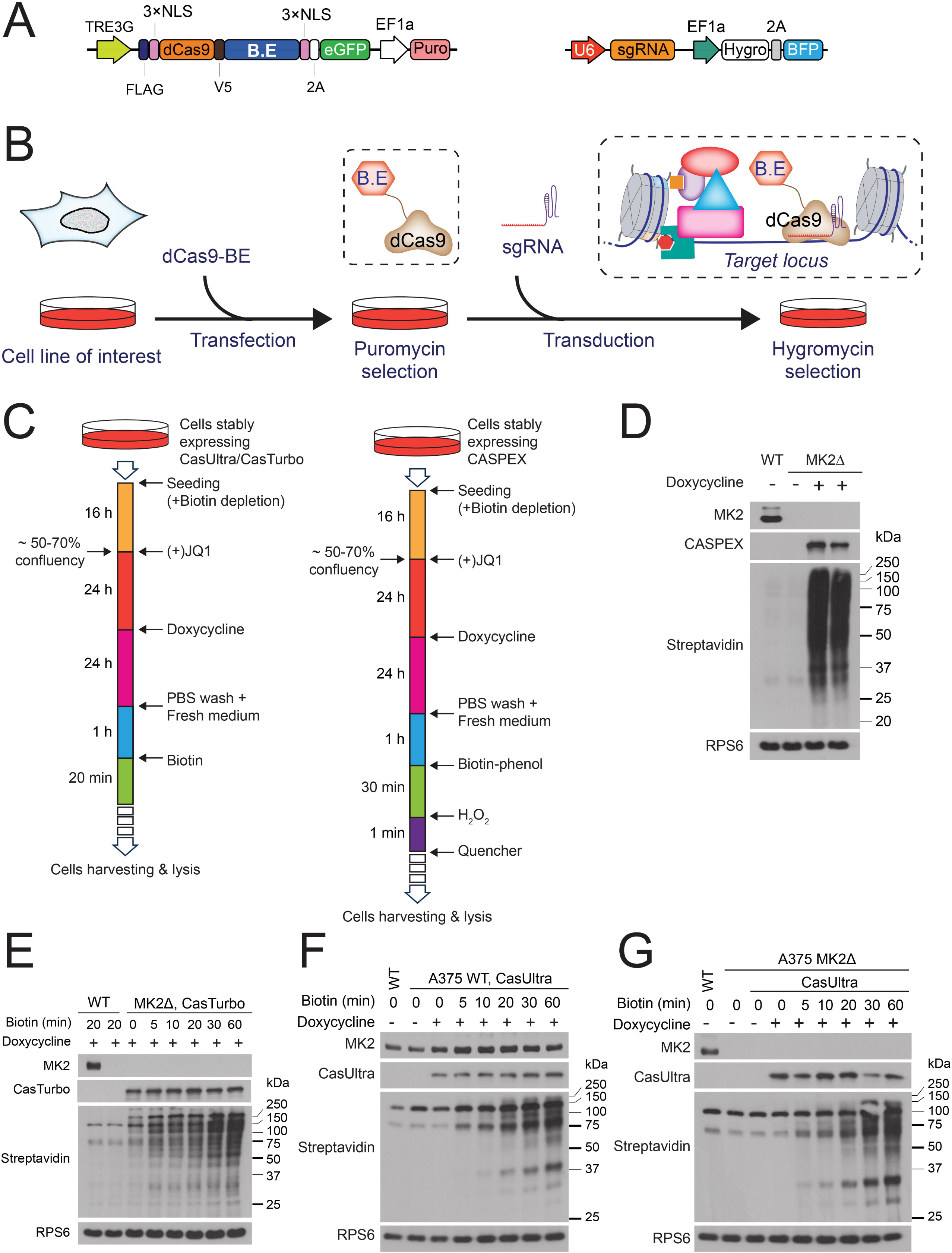
Generation and characterization of dCas9-BE models. **A.** Schematics of the lentiviral sgRNA (left) and inducible dCa9-BE (right) constructs. **B.** Workflow of dCas9-BE cell line generation. **C.** Experimental workflow for CasTurbo/CasUltra (left) and CASPEX (right). **D.** WB analysis of CASPEX expression and biotinylation ability in A375 lines. Biotinylation time: 1 min. **E–G.** WB analyses of the biotinylation kinetics of CasTurbo (**E**) and CasUltra in WT A375 cells (**F**) and MK2Δ A375 cells (**G**). Biotinylation was triggered by adding 500 µM biotin for the indicated times. Biotinylated proteins were detected with HRP-streptavidin. Results are representative of two biological replicates. RPS6 was used as a loading control.

Primary characterization by WB analysis and fluorescence microscopy imaging were performed after inducing the dCas9-BE and the sgRNA. Following dCas9-BE transfection and selection, we validated the inducible expression, expression level, cellular distribution, and biotinylation ability. Clones with moderate dCas9-BE expression that localized exclusively in the nucleus were selected for WB and IF analyses. After inducing dCas9-BE expression and biotinylation (**Figure 2C),** the samples were processed for WB to assess expression and biotinylation. CASPEX resulted in a high level of biotinylation after a 1-min reaction time (**Figure 2D**), while CasTurbo (**Figure 2E**) and CasUltra (**Figure 2F-G**) displayed a time-dependent increases in biotinylation intensity over the first 30 min. The dCas9-BE transcript is fused to GFP, while the sgRNA construct also expresses BFP, allowing for fluorescent imaging and cell sorting. Populations with stable co-expression of dCas9-BE and the sgRNA were characterized for their dCas9-BE inducibility, localization, and biotinylation ability by WB and imaging analyses, then sorted to achieve similar levels of dCas9-BE/sgRNA co-expression and reduce variations between biological samples and different passage numbers. Briefly, cells were first sorted based on their BFP expression, and the cells with high BFP expression were sorted into low- and high-GFP-expressing cell populations (**Figure 3A**). Live cell imaging revealed that both the CasUltra LentiG and 053MYC-expressing lines were constitutively and uniformly BFP-positive and inducibly GFP-positive, verifying the sorting efficiency of the different models (**Figure 3B**).

**Fig. 3:**
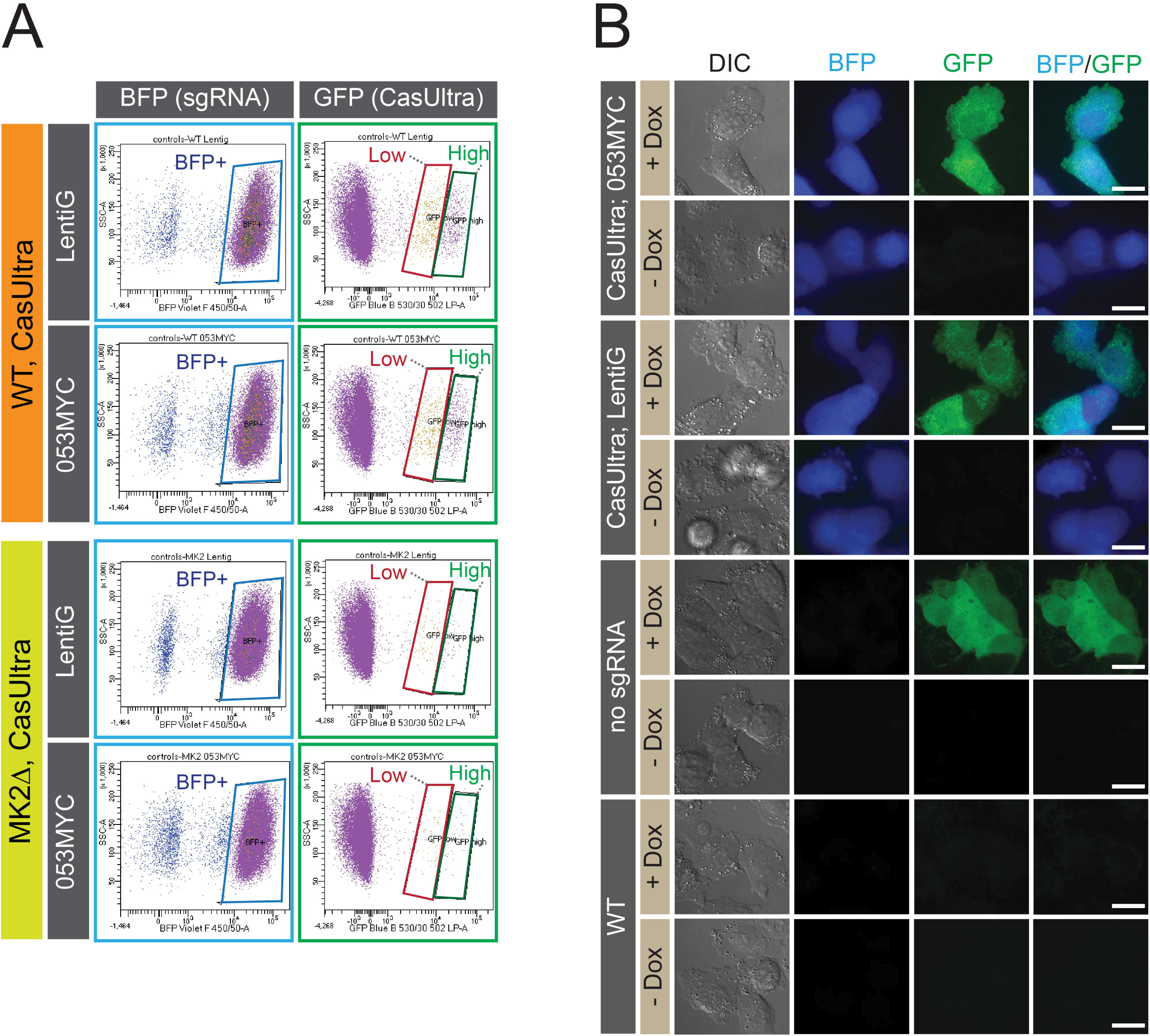
CasUltra models sorting and characterization. **A.** Scatter plots of sorted WT (top) and MK2Δ (bottom) co-expressing sgRNA and CasUltra constructs based on BFP and GFP fluorescence levels. **B.** Live-cell imaging of FACS-sorted cell populations with high expression of CasUltra. The WT line was used as a BFP- and GFP-negative control. Scale bars: 10 µm.

### Expressing dCas9-BE does not affect BET protein inhibitor sensitivity

We next examined if targeting dCas9-BE to the *MYC* promoter affected proliferation rates by disrupting the local chromatin proteome. MTT assays of cells co-expressing CasTurbo with either sg053MYC or sgNS (as a negative control) did not reveal significant impacts on proliferation under basal conditions (**Figure 4A**). We next tested whether dCas9-BE expression was likely to modify the cellular response to BETi. Crystal violet-based proliferation assays of cells co-expressing CasTurbo with either sg053MYC or sgNS under various BET inhibitors did not reveal changes in proliferation (**Figure 4B**).

**Fig. 4:**
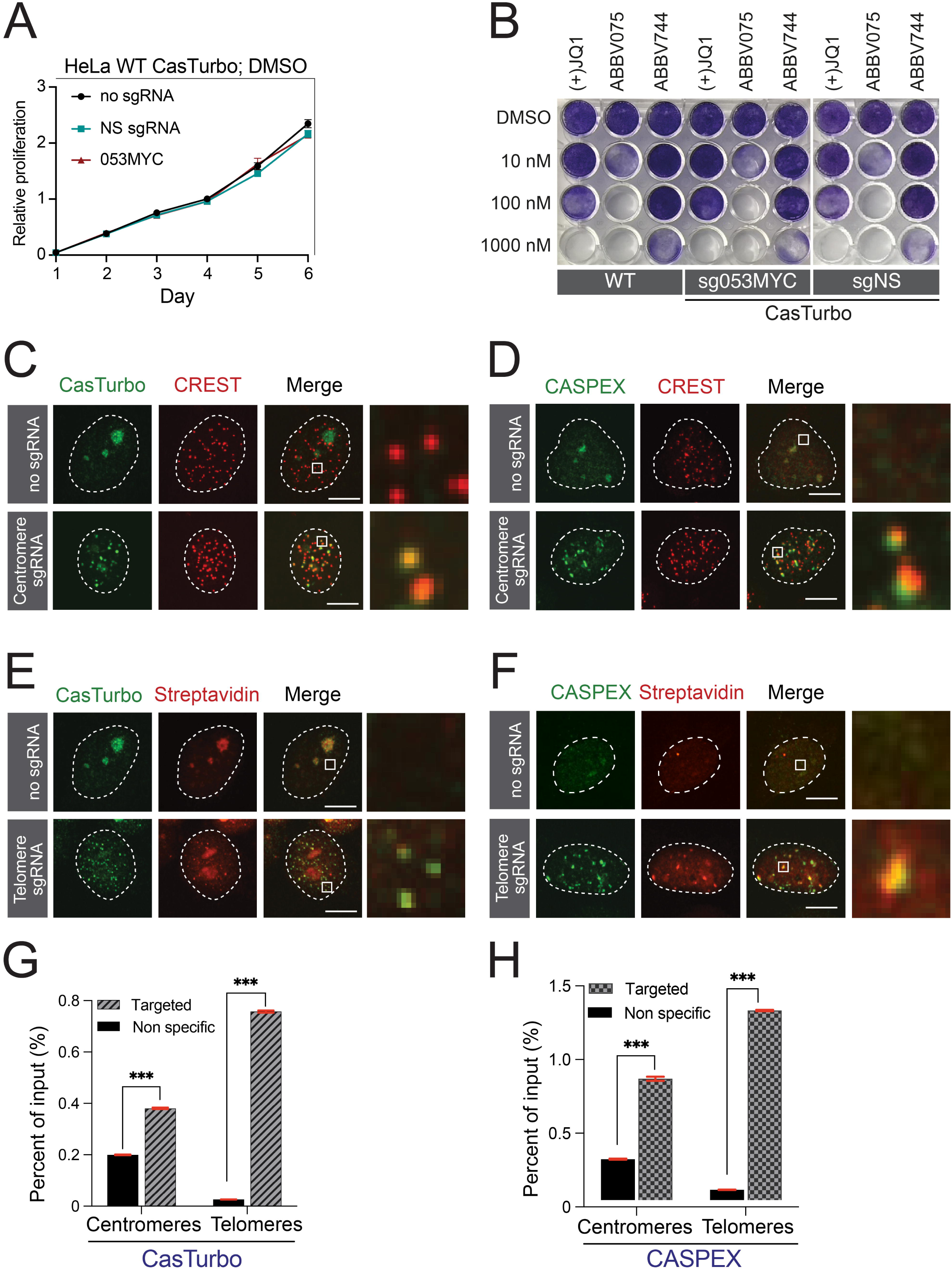
Targeting telomeres and centromeres using CASPEX and CasTurbo. **A.** Proliferation assays of HeLa cells co-expressing CasTurbo and sg053MYC, sgNS, or no sgRNA over 6 d. **B.** Clonogenic assays of HeLa cells co-expressing CasTurbo, or not, and sg053MYC, sgNS, or no gRNA and treated with increasing concentrations of the BET inhibitors JQ1, ABBV075, and ABBV774 for 6 d. **C–F**. IF of HeLa cells co-expressing CasTurbo or CASPEX and either no sgRNA or an sgRNA targeting the centromeres or telomeres. Representative images from two independent experiments are shown. Scale bars: 10 µm. **G, H.** FLAG ChIP-qPCR to detect 3×FLAG-tagged CasTurbo (**G**) and CASPEX (**H**) at the telomeres and centromeres. Results are expressed as the percent enrichment compared to the input sample. Data are the mean ± SD. *** *P* < 0.001, by *t*-test.

### Targeting CasTurbo to the centromeres and telomeres

To ensure that the dCas9-BEs localized to their targets properly, we first targeted telomeres and centromere alpha-satellites—both regions with repetitive sequences—as a proof-of-concept. FACS-sorted cell populations co-expressing CasTurbo or CASPEX and the alpha satellite-targeted sgRNA revealed colocalization between the dCas9-Bes and the centromere marker CREST (**Figure 4C-D**). In contrast, the dCas9-BEs had diffuse and predominantly nucleolar distributions in the absence of the sgRNA. Next, we verified the biotinylation efficiency of the targeted loci by IF imaging of the biotinylated proteins. The biotinylation spots overlapped with the distribution of dCas9-BE, indicating efficient biotinylation at the targeted loci (**Figure 4E-F**). Lastly, we assayed the locus-targeting accuracy of our dCas9-BE systems using ChIP-qPCR. To accomplish this, we used sgNS as a negative control for the telomere- and centromere-targeting sgRNAs. As expected, dCas9-BE and the biotinylation proteins were significantly enriched at the target locus compared with the negative control (**Figure 4G-H**).

### CasTurbo-based proteomic profiling of telomeres and centromeres

The CasTurbo sgCentromere and sgTelomere models were processed in tandem with two control lines, expressing either sgNS or no sgRNA. Samples were biotinylated as per **Figure 2C** without JQ1 treatment. Biotinylated proteins were purified and digested, and the peptides were isopropanol-fractionated then analysed by MS. To assess the quality of the CasTurbo-based MS dataset, we first evaluated the overall protein enrichment, which revealed similar total protein content across the samples (**Figure 5A**). Principal component analysis (PCA) clustered the samples based on their sgRNAs, as expected (**Figure 5B**). CasTurbo co-expressed with sgTelomere identified 1105 proteins. Gene ontology (GO) analysis highlighted 19 proteins with known telomerase and telomeric DNA-binding activities, including 5/6 subunits of shelterin (TERF1, TERF2, ACD, TINF2, and POT1), a telomere-binding protein complex (64), and numerous additional telomere-interacting proteins, such as TERF2IP, the bromodomain proteins BRD9 and BRWD1, and the histone deacetylase HDAC3 (**Figure 5C**). CasTurbo co-expressed with sgCentromere identified 1189 proteins, including known centromeric proteins like CENPT, CENPU, CENPC, and INCENP (**Figure 5C**) and centromere-related hits, including BRCA1/BRCA2 (65), IPO5, CUX1, and ZNF451, which were previously validated as CENPA-interactors (66). Despite these very encouraging results obtained with CasTurbo, we observed a significant level of non-specific biotinylation. This was particularly linked to the tendency of CasTurbo to accumulate in nucleoli because of TurboID’s DNA-binding activity.

**Fig. 5:**
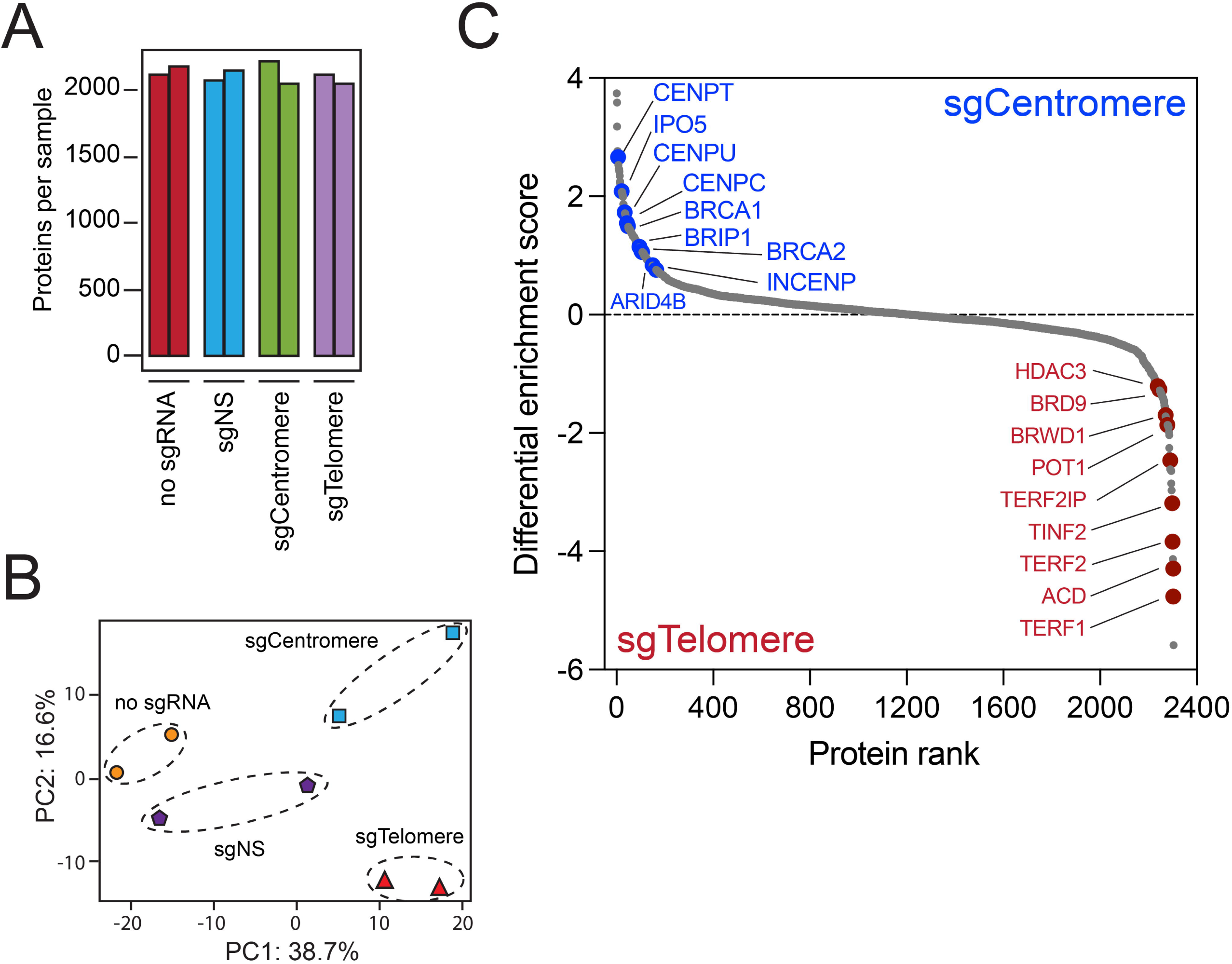
MS-based profiling of telomeres and centromeres with CasTurbo. **A.** Total proteins quantified across the CasTurbo samples. **B.** PCA plot of CasTurbo-enriched proteins. **C.** Protein rank plot of differentially enriched proteins between the sgCentromere- and sgTelomere-expressing samples.

### BET inhibition modulates the chromatin composition at the *MYC* promoter

As the CasTurbo approach was effective at repetitive sequences, we sought to sharpen its focus by applying it at the non-amplified *MYC* locus. To avoid the nucleolar localization observed with TurboID (**Figure 4C-D**), we used CasUltra (using the smaller UltraID, which lacks a DNA binding domain (47) (**Figure 6A**)) to profile the changes induced by JQ1 at the *MYC* promoter in WT and MK2 KO A375 melanoma cells. CasUltra cell lines with high GFP expression were processed for MS analysis as in **Figure 6B-C**. Given the low number of loci being targeted, we fractionated the peptides prior to MS analysis as in **Figure 6D**. Approximately 2000 proteins were quantified in each CasUltra sample (**Figure 7A**), which clustered according to their JQ1 treatment and MK2 expression (**Figure 7B**). Following JQ1 treatment, the *MYC* promoter environment of WT cells was enriched with proteins associated with GTPase, GDP/GTP binding, and nucleoside triphosphate activities (**Figure 7C-D**). The *MYC* promoter of MK2 KO cells were also enriched for proteins with nucleoside triphosphate activity but remained enriched with proteins associated with RNA polymerase II even after JQ1 treatment (**Figure 7E-F**). Sustained *MYC* transcription in MK2 KO cells following JQ1 treatment is consistent with the BET inhibitor resistance we recently reported (50). On the other hand, the profiles of proteins displaced from the *MYC* promoter in DMSO- and JQ1-treated WT and MK2 KO cells were similar (**Figure 8A-D**). Together, these results suggest that the loss of MK2 enhances resistance to BETi by retaining key chromatin regulators at specific loci, such as *MYC* (**Figure 8E-F**).

**Fig. 6:**
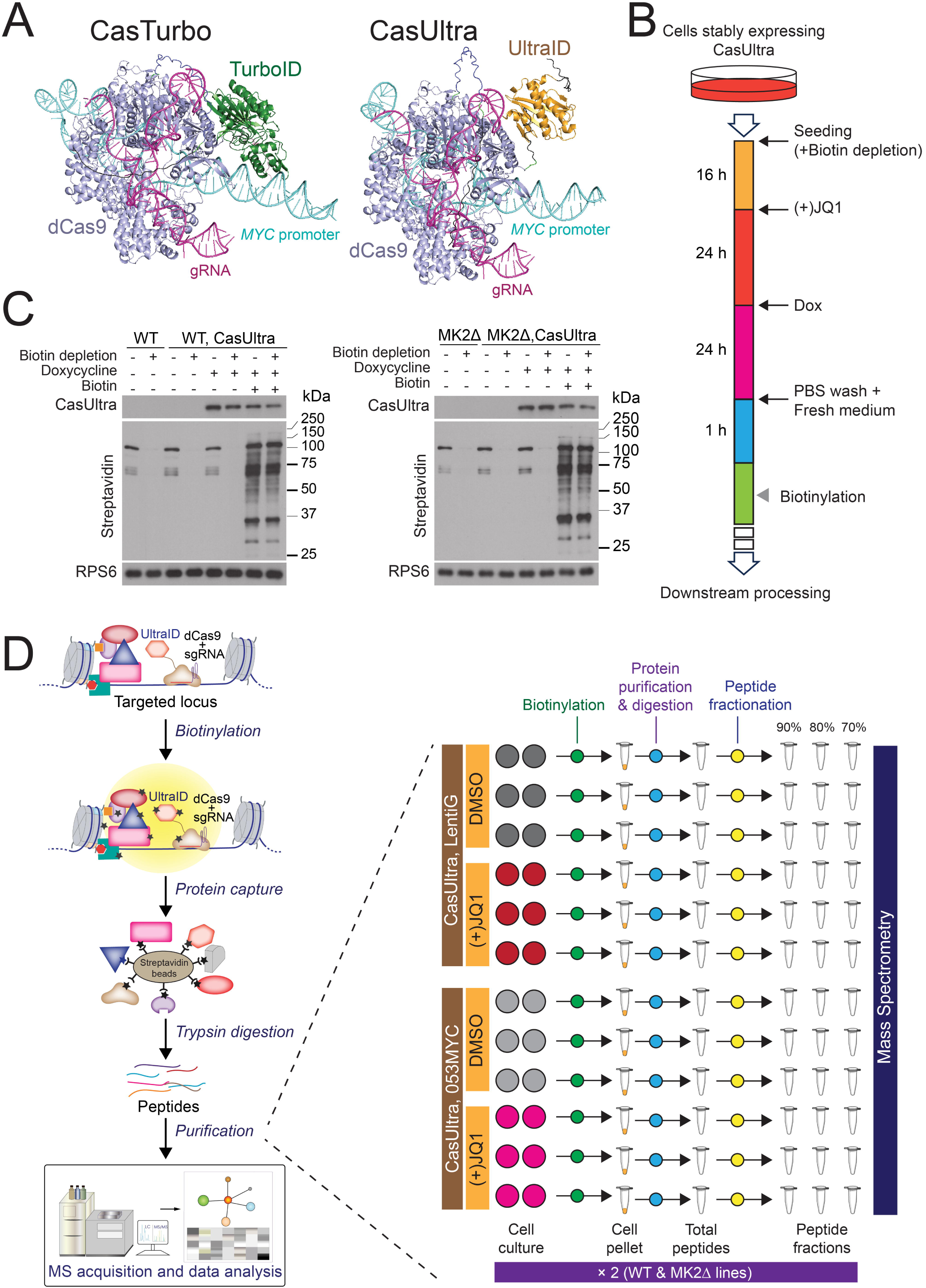
Experimental design for MS-based CasUltra analysis. **A.** Structural modelling of the CasTurbo (left) and CasUltra (right) constructs in a complex containing the sgRNA and the *MYC* promoter. The sequences used for AlphaFold3 modelling are available in **Supplementary Table S1C**. **B.** The CasUltra-based workflow for MS analysis. **C.** WB analysis of WT (left) and MK2Δ (right) A375 cells prepared for CasUltra proteomics with and without biotin depletion. **D.** The CasUltra workflow for locus-targeted MS-based proteomics (left). Schematic of the experimental design for MS analysis (right).

**Fig. 7:**
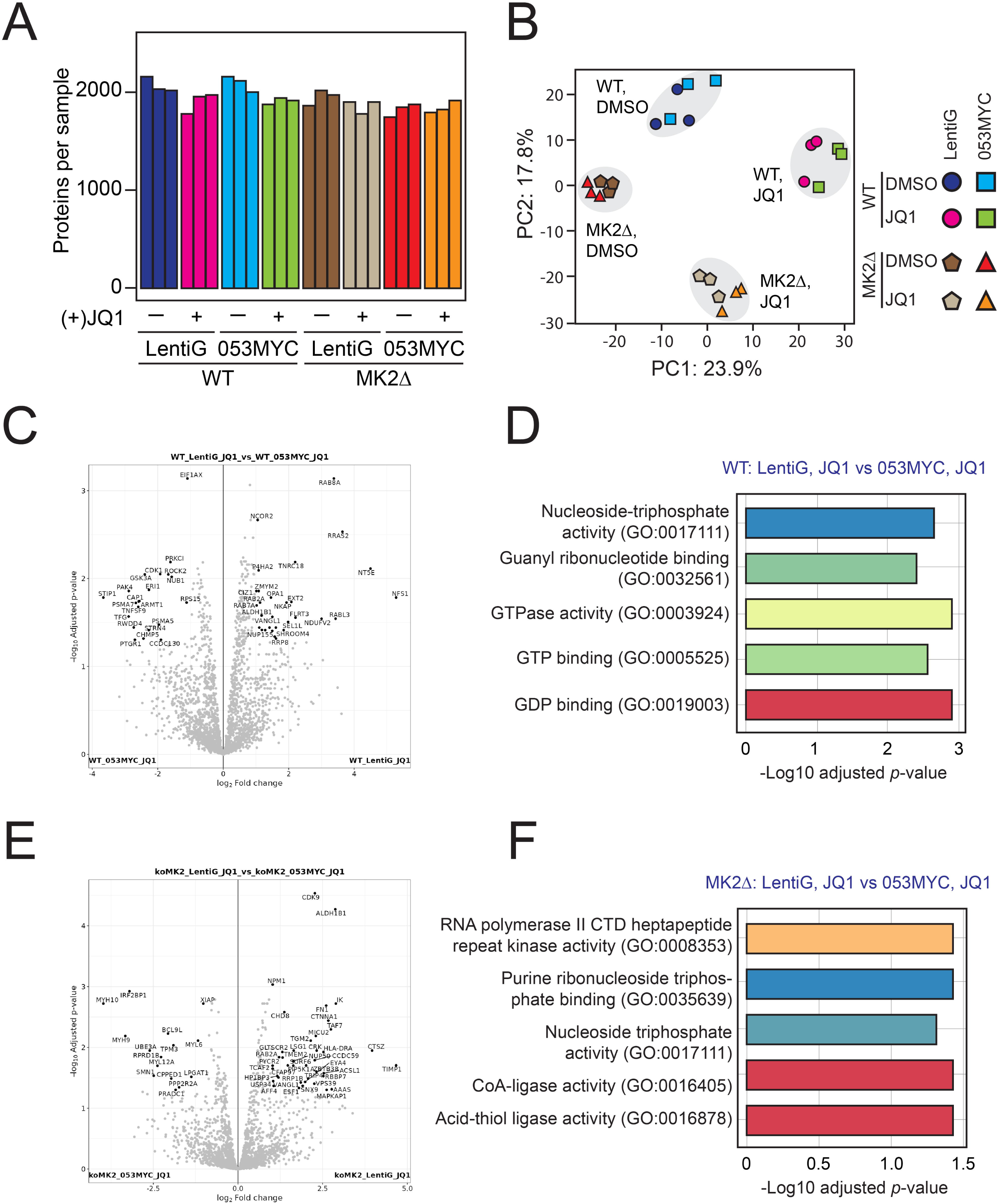
MS-based elucidation of the *MYC* promoter protein environment following JQ1 treatment. **A.** Total proteins quantified across the CasUltra samples. **B.** PCA plot of MS data from CasUltra-enriched proteomes. **C-F.** Volcano plots representing differential enrichment of biotinylated proteins at the *MYC* promoter and top enriched GO molecular function terms in the 053MYC-targeting CasUltra proteome compared with an empty LentiGuide control following JQ1 treatment in WT (**C-D**) and MK2Δ (**E-F**) cell lines. Proteins are represented by dots. Black labelled dots represent positively enriched proteins at significance thresholds of adjusted *p*-value = 0.05 and log_2_ fold change cut-off = 1.

**Fig. 8:**
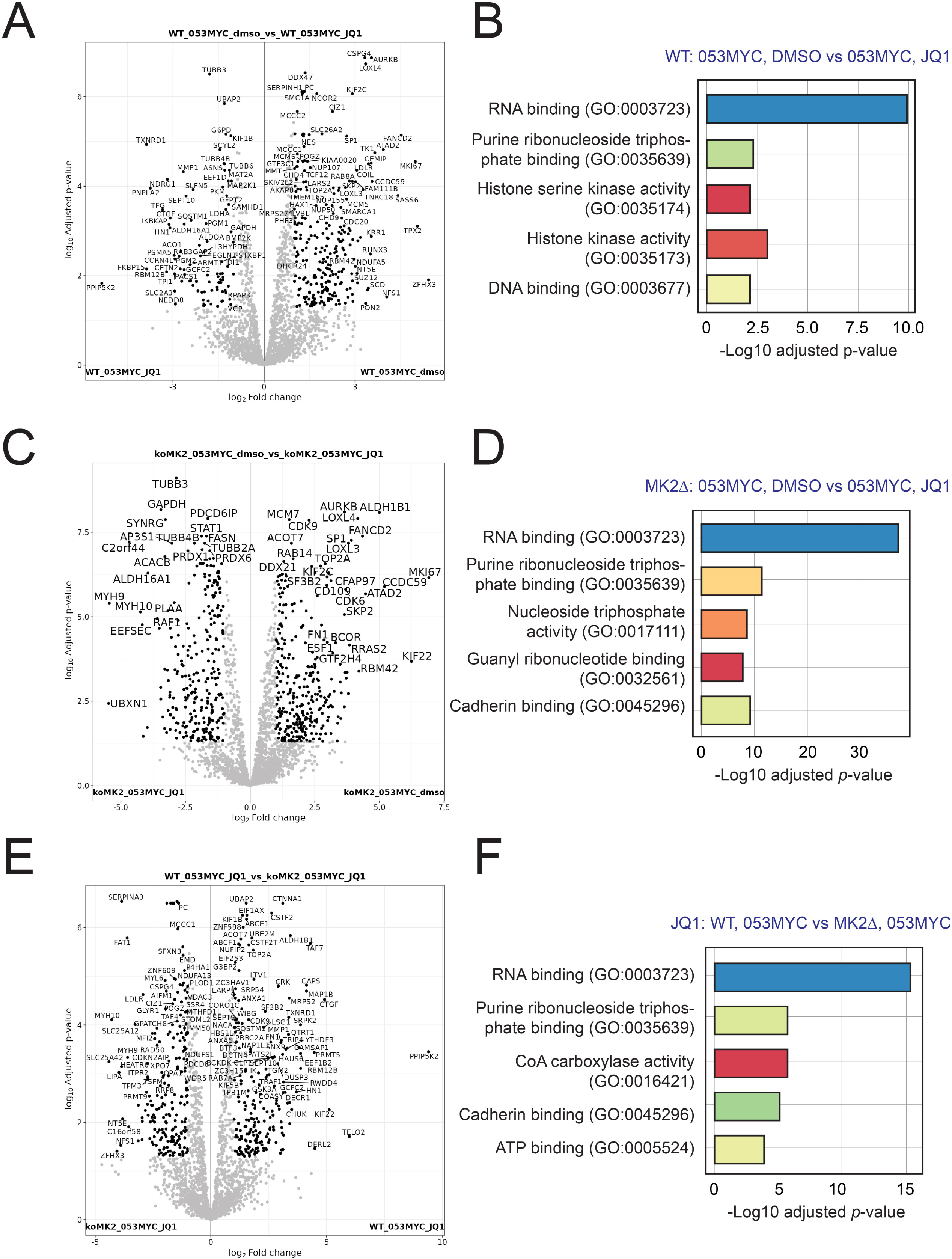
Effect of MK2 KO on *MYC* promoter reorganization following JQ1 treatment A-D. Volcano plots representing differential enrichment of biotinylated proteins at the *MYC* promoter and top enriched GO molecular function terms induced by JQ1 treatment compared to DMSO in WT (71) and MK2Δ (**C-D**) lines. **E-F.** Volcano plot (**E**) and GO term analysis (**F**) of differences in the *MYC* promoter proteomes of WT and MK2Δ cells following JQ1 treatment.

## Discussion

Locus-specific proteomic tools are a promising means to determine the molecular mechanisms underlying the organization, use, repair, and protection of genetic information. Among these tools, the initial development of the dCas9-BE approach has sparked much hope and enthusiasm in the chromatin science community despite the significant hurdles to overcome. Here, we described CasTurbo and CasUltra, two new dCas9-BE tools that allow for rapid profiling of locus-specific proteomes in live cells under mild conditions. As a proof of concept, we used CasTurbo to identify the proteins associated with centromeres and telomeres. However, we observed that the CasTurbo constructs accumulated at the nucleolus in addition to their genomic target, leading to the purification of nucleolar proteins such as TCOF1. We hypothesized that the N-terminal domain of TurboID, which contains a DNA-binding site, contributed to the non-specific chromatin distribution of CasTurbo. As the recently described UltraID abortive biotin ligase lacks the equivalent N-terminal sequence (47), we selected it to characterize the *MYC* promoter. JQ1 has been reported to downregulate the expression of several genes involved in tumour maintenance and progression, including *MYC* (5, 26). The JQ1-induced disturbance in the *MYC* promoter’s proteome reduces its expression and limits cellular self-renewal, making BET inhibitors good therapeutic candidates for MYC-driven cancers such as multiple myeloma (26, 67, 68). CasUltra-based *MYC* promoter proteomics enabled the identification of numerous proteins known to interact with the regulatory elements governing its transcription, such as BRD9 (69), ARID2 (70), and co-regulators orchestrating its expression dynamics. Our comparative analysis of proteomic data across various experimental conditions reinforces our previous observation that MK2 ablation reduces the remodelling of chromatin complexes following JQ1 treatment (50). For instance, we observed decreases in the subunits of DNA remodelling complexes, including PBRM1; ARID2; SMARCC2; and numerous other BRD-containing proteins, such as BAZ1B and ATAD2. The CasUltra approach has enhanced our understanding of chromatin composition following loss of MK2. For example, deleting MK2 enhanced the recruitment of numerous chromatin proteins at the *MYC* promoter, including cyclin-dependent kinases (CDK6 and CDK9), chromatin remodellers (ATAD2, PBRM1, SMARCC2, and BAZ1B), and transcription factors (ZNF24 and ZNF326). Even after JQ1 treatment, MK2 KO cells displayed increased enrichment in transcription initiation factors (TAF4), chromatin remodeller SWI/SNF subunits (SMARCA1 and SMARCA2), DNA-repair proteins (RAD50), mediator complex subunits (MED24), and ZNF transcription factors (ZNF609).

In summary, we have expanded the toolbox for locus-specific proteomics by developing CasTurbo and CasUltra, which enable rapid and gentle proteomic mapping of specific genomic loci in living cells. We found that the rapid biotinylation provided by CasTurbo was counter-balanced by its non-specific distribution, making CasUltra the best dCas9-BE construct to map dynamic protein complexes at the genomic locus level. Notably, the spatial and temporal organization of protein complexes on the chromatin will reveal insights into how other signalling pathways function at the single genomic locus level, which remains a challenging task.

## Acknowledgments

The pSpCas9(BB)-2A-Puro (PX459) V2.0 plasmid (Addgene #62988) was donated by Feng Zhang, the inducible CASPEX expression plasmid (Addgene #97421) was gifted by Steven Carr and Samuel Myers, and the lentiGuide-Hygro-mTagBFP2 plasmid (Addgene #99374) was provided by Kristen Brennand. Research in the Lambert laboratory is funded by a Discovery Grant from the Natural Sciences and Engineering Research Council of Canada (NSERC; 1304616-2017), an Operating Grant from the Cancer Research Society (CRS; 935296), and Leader’s Opportunity Funds from the Canada Foundation for Innovation (37454). J.-P.L. was supported by a Junior 2 salary award from the Fonds de Recherche du Québec-Santé (FRQS). P.-E.K.T. was supported by a Bourses de formation Desjardins pour la recherche et l’innovation from the Fondation du CHU de Québec, a Bourse Distinction Luc Bélanger from the Cancer Research Centre – Université Laval, and a FRQS doctoral training scholarship. Funding sources were not involved in the study design, data collection, analysis, and interpretation of data, and in the writing of the manuscript.

## Data Availability

All MS files generated as part of this study have been deposited to the MassIVE repository. Cell lines and plasmids created or employed in this study will be provided upon request unless created by others or commercially available.

## Conflicts of interest

none

## Author contributions

Conceptualization, J.P.L; Methodology, P.E.K.T. and J.P.L; Software, J.P.L.; Investigation, P.E.K.T., J.L., and J.P.L.; Data Curation, J.P.L.; Writing – Original draft, P.E.K.T. and J.P.L; Writing – Review and Editing, P.E.K.T. and J.P.L.; Supervision, J.P.L.; Project Administration, J.P.L.; and Funding Acquisition, J.P.L.

## Supplemental data

This article contains supplemental data.

## Abbreviations

BET: Bromodomain and extra-terminal
BETi: BET inhibition
BRD: Bromodomain
CASPEX: Catalytically dead Cas9 (dCas9) fused to the biotinylation enzyme APEX2
CasTurbo: Catalytically dead Cas9 (dCas9) fused to the engineered biotin ligase TurboID
Casula: Catalytically dead Cas9 (dCas9) fused to the engineered biotin ligase UltraID
ChIP: Chromatin immunoprecipitation
dCas9-BE: Catalytically dead Cas9 (dCas9) fused to a biotinylation enzyme
Kac: Acetylated lysine
LC: Liquid chromatography
MS: Mass spectrometry

## Notes

### Competing Interest Statement

The authors have declared no competing interest.

